# Altered cerebellar response to somatosensory stimuli in the *Cntnap2* mouse model of autism

**DOI:** 10.1101/2021.05.13.443762

**Authors:** Marta Fernández, Carlos A. Sánchez-León, Javier Llorente, Teresa Sierra-Arregui, Shira Knafo, Javier Márquez-Ruiz, Olga Peñagarikano

## Abstract

Atypical sensory processing is currently included within the diagnostic criteria of autism. The cerebellum is known to integrate sensory inputs of different modalities through its connectivity to the cerebral cortex. Interestingly, cerebellar malformations are among the most replicated features found in postmortem brain of individuals with autism. We studied cerebellar integration of sensory information in a mouse model of autism, knockout for the *Cntnap2* gene. *Cntnap2* is widely expressed in Purkinje cells and has been recently reported to regulate their morphology. Further, individuals with *CNTNAP2* mutations display cerebellar malformations and CNTNAP2 antibodies are associated with a mild form of cerebellar ataxia. Previous studies in the *Cntnap2* mouse model show an altered cerebellar sensory learning. However, a physiological analysis of cerebellar function has not been performed yet. We studied sensory evoked potentials in cerebellar Crus I/II region upon electrical stimulation of the whisker pad in alert mice and found striking differences between WT and *Cntnap2* KO mice. In addition, single-cell recordings identified alterations in both sensory-evoked and spontaneous firing patterns of Purkinje cells. These changes were accompanied by altered intrinsic properties and morphological features of these neurons. Together, these results indicate that the *Cntnap2* mouse model could provide novel insight into the pathophysiological mechanisms of ASD core sensory deficits.

## Introduction

Atypical sensory processing is currently included within the diagnostic criteria of autism (APA, 2013). The cerebellum is known to play a role in integration of different sensory modalities (e.g. hearing, sight, touch and smell) through its connectivity to the cerebral cortex (Proville et al., 2014). Interestingly, cerebellar malformations are among the most replicated features found in postmortem brain of individuals with autism (S. S. Wang, Kloth, & Badura, 2014) and alterations in functional connectivity between the cerebellum and cortical sensory areas have been found through fMRI (Lidstone, Rochowiak, Mostofsky, & Nebel, 2021).

Loss of function mutations in the *CNTNAP2* gene are associated with a syndromic form of autism that presents with cerebellar abnormalities, including hypoplasia of the cerebellar vermis and hemispheres (Rodenas-Cuadrado et al., 2016). Further, autism-linked common genetic variation in *CNTNAP2* has been associated with reduction in cerebellar grey matter volume, as determined by MRI (Tan, Doke, Ashburner, Wood, & Frackowiak, 2010). Incidentally, CNTNAP2 antibodies have been identified in sera from patients with otherwise unexplained progressive cerebellar ataxia with mild to severe cerebellar atrophy (Becker et al., 2012; Melzer, Golombeck, Gross, Meuth, & Wiendl, 2012), supporting the role of this gene in cerebellar development and function. *CNTNAP2* is widely expressed in the cerebellum (Gordon et al., 2016), and it has been recently reported to regulate Purkinje cell (PC) morphology (Argent et al., 2020). In mice, loss of *Cntnap2* function causes autism-like behaviors (Penagarikano et al., 2011), as well as several sensory abnormalities such as hypersensitivity to painful stimuli (Dawes et al., 2018) and alterations in auditory (Scott et al., 2018) as well as olfactory behaviors (Gordon et al., 2016). Neuroanatomical analysis of this model shows an alteration in cerebellar volume, as measured by structural MRI (Ellegood & Crawley, 2015). Classical tests measuring cerebellar function report improved performance in the accelerating rotarod (Penagarikano et al., 2011), unstable gait (Argent et al., 2020) and cerebellar sensory learning defects (Kloth et al., 2015). However, a physiological analysis of cerebellar function has not been performed yet. In this work, we studied cerebellar integration of sensory information in alert *Cntnap2* mice. We discovered alterations in the firing patterns of PC both spontaneous as well as in the evoked response to sensory stimuli. This alteration was accompanied by an increased excitability of PC and reduced dendritic complexity in these neurons. Together, these results provide novel insight into the pathophysiological mechanisms by which CNTNAP2 mutations cause impairments in cerebellar function that may contribute to ASD core deficits.

## Results

### Spontaneous *in vivo* Purkinje cell activity is altered in *Cntnap2*^-/-^ mice

We first performed extracellular recordings of spontaneous PC activity in the Crus I/II area of awake animals (**Figure 1A**). Crus I/II was selected because of its functional relationship with sensory whisker inputs (Gao et al., 1996) and its association with ASD (D’Mello, Moore, Crocetti, Mostofsky, & Stoodley, 2016; Skefos et al., 2014; Y. Wang, Xu, Zuo, Zhao, & Hao, 2020). PC present two types of firing patterns: simple spikes (SS) and complex spikes (CS), generated by distinct inputs (parallel fibers and climbing fibers, respectively) and distinguished by their particular waveforms. Despite their different origin interactions between the two types of spikes have been described, such that CS firing is thought to modulate SS activity (Tang et al., 2017). We found a lower CS firing frequency in *Cntnap2* KO mice compared with WT controls (**Figure 1B**). The firing frequency of SS (**Figure 1C**), as well as the predominant, or preferred, SS firing frequency (**Figure 1D**) were not different between genotypes. However, *Cntnap2* KO mice show a higher irregularity in the temporal firing pattern of SS (**Figure 1E**), as measured by the coefficient of variation (CV) of the inter-spike intervals. Such difference is not observed when adjacent inter-spike intervals of presumably burst firing are considered, denoted by CV_2_ (**Figure 1F**).

**Figure 1.**
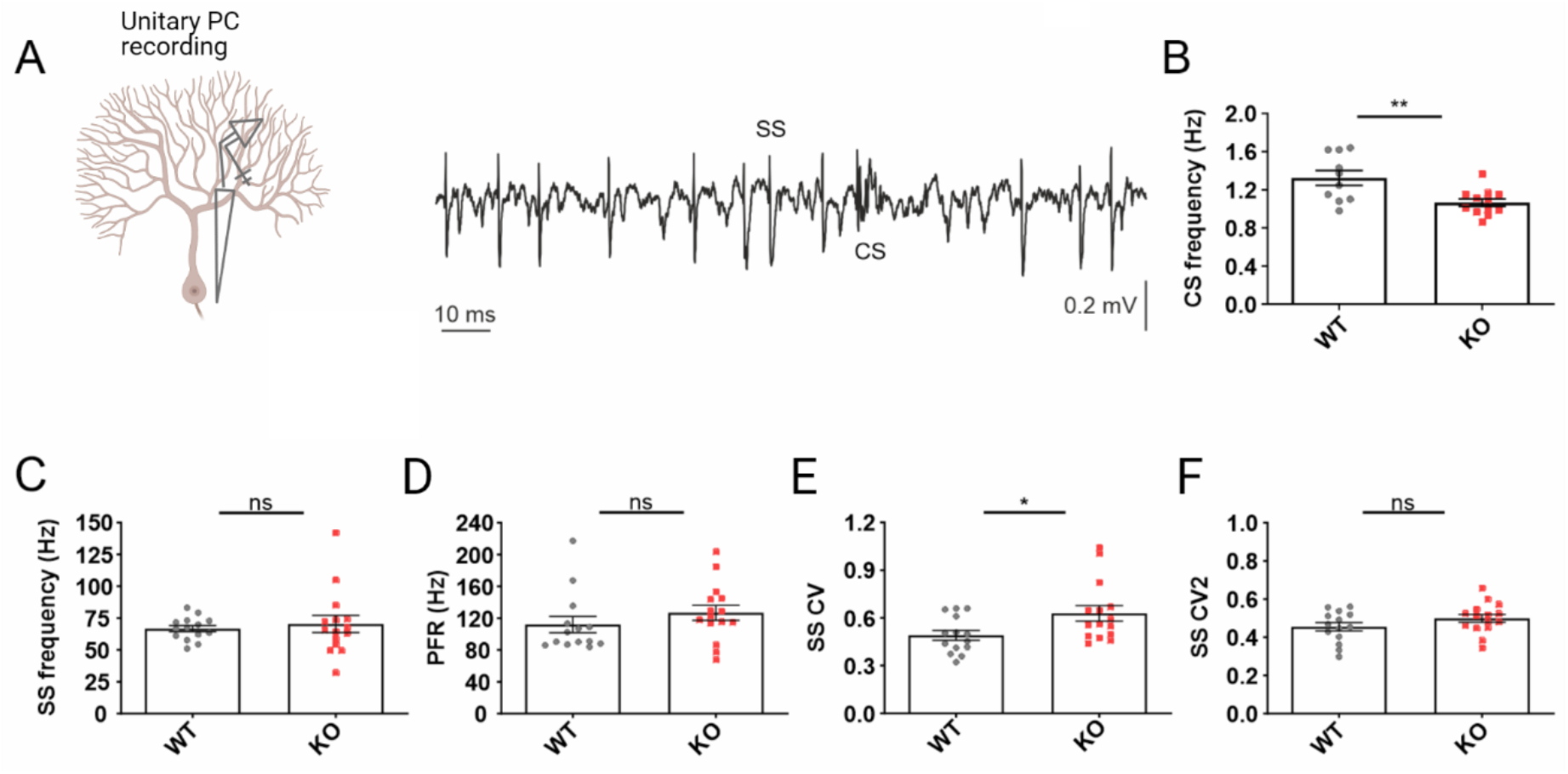
*In vivo* spontaneous activity of Purkinje cells. (A) Schematic illustration of an extracellular PC recording and a representative trace displaying simple spikes (SS) and complex spikes (CS). (B) CS firing rate. (C) SS firing rate. (D) SS predominant firing rate. (E) Coefficient of variation (CV) of the inter-spike intervals for SS. (F) Coefficient of variation for adjacent inter-spike intervals (CV_2_). Data are presented as mean ± S.E.M. n=14-15 neurons/genotype. Student’s t-test (*p<0.05, **p<0.01).

### Purkinje cell responses to sensory-evoked stimuli are altered in *Cntnap2*^-/-^ mice

To characterize PC activity in response to somatosensory stimuli, we recorded the Local Field Potential (LFP) near the PC layer from the cerebellar Crus I/II area after subcutaneous electrical stimulation of the ipsilateral whisker pad in alert mice (**Figure 2A**).This protocol induces a more reproducible response than tactile whisker stimulation (Marquez-Ruiz & Cheron, 2012).

**Figure 2.**
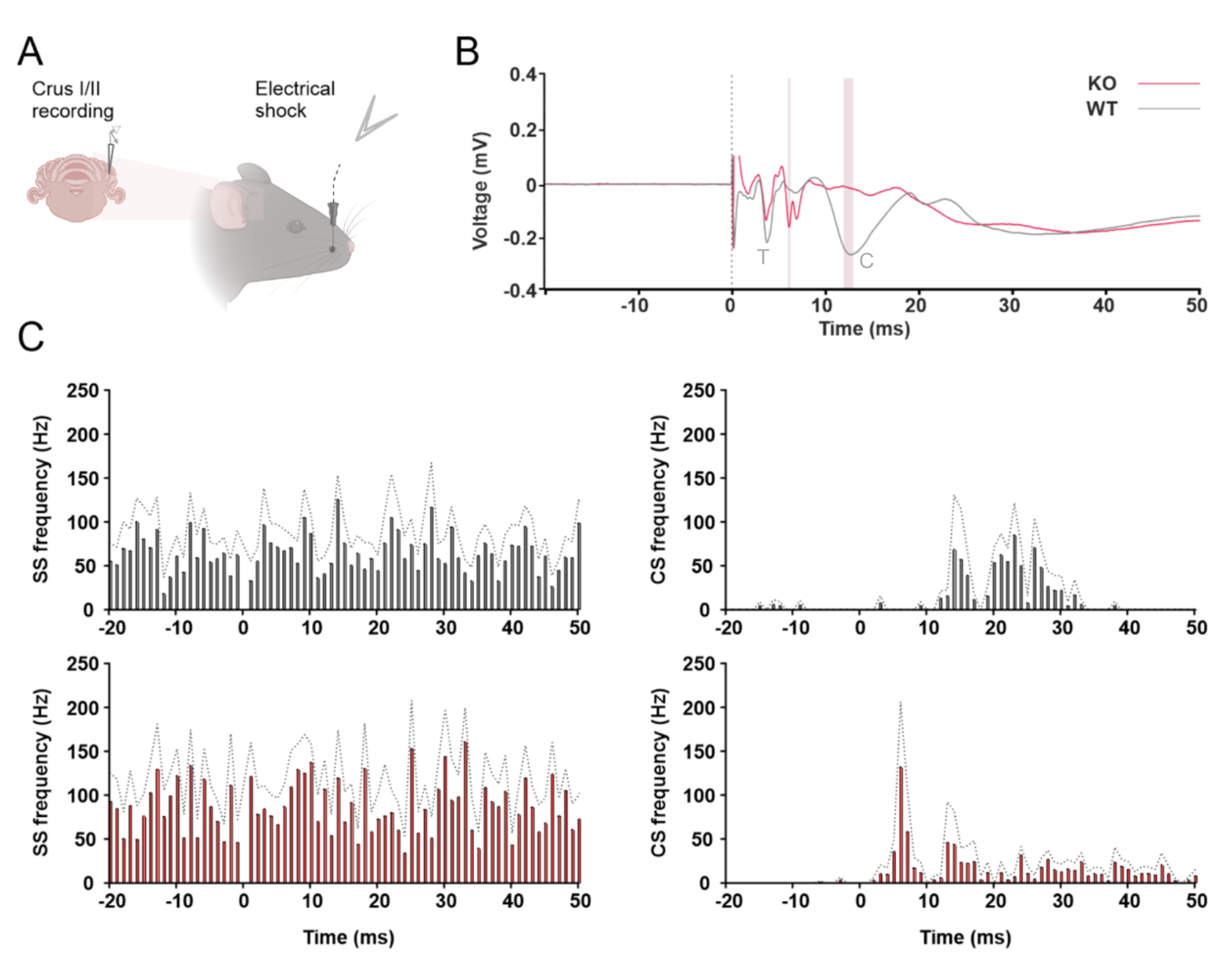
Cerebellar LFP after electrical stimulation of the whisker pad. (A) Schematic illustration representing the LFP recording. (B) Event-related potential (ERP) analysis comparing the average SEP traces for 11 KO (red trace) and 9 WT mice (black trace). Vertical bars indicate statistically different latencies, corresponding to the cortical peak (C) in WT at 12.87±0.40 ms, which is absent in KOs, who show a novel negative peak, presumably an anticipated cortical response, at 6.46±0.14 ms. Student’s t-test (p<0.05). (C) Temporal firing pattern for SS (left) and CS (right) in WT (black) and KO (red) mice after electrical stimulation. Note that CS appear at post-stimulation latencies concordant with the C component in WT, while they appear earlier in KO mice, likely indicating an anticipated C response in the animals. The graphs represent the mean ± S.E.M (discontinuous line). n=7-10 PC per genotype.

Concordant with what was previously described (Marquez-Ruiz & Cheron, 2012), in wild-type (WT) mice this electrical stimulation evoked a highly reproducible sensory evoked potential (SEP) with two main negative components appearing at around 3 and 12 ms (3.94±0.13 ms and 12.87±0.40 ms), corresponding to trigeminal (T) and cortical (C) pathways, respectively. Strikingly, the waveform of SEP in *Cntnap2* knockout (KO) mice was notably different, showing the expected T wave (4.10±0.15 ms) and a novel negative peak at around 6ms (6.46±0.14 ms) in the absence of the expected C wave at 12 ms (**Figure 2 B**). To determine the basis for the altered SEP response observed in *Cntnap2*^-/-^ mice, we then recorded unitary PC upon whisker stimulation. Previous studies have correlated the appearance of a SS burst with the above-mentioned T component, whereas CS occurred at post-stimulation latencies concordant with the C component (Marquez-Ruiz & Cheron, 2012; Mostofi, Holtzman, Grout, Yeo, & Edgley, 2010). In agreement with this, in WT mice we observed CS appearing 12.46 ms after the stimulus onset, coinciding with the C wave observed in the SEP. In *Cntnap2*^-/-^ mice, however, the appearance of the CS takes place at 6 ms, matching the latency of the novel peak observed in the corresponding SEP (**Figure 2C**). Although the precise origin of this novel peak needs to be elucidated, the data suggest that the cerebellar representation of cortical input is anticipated in *Cntnap2*^-/-^ mice.

### Increased Purkinje cell intrinsic excitability in *Cntnap2* KO mice

To assess if alterations in the intrinsic properties of PC could be responsible for the aberrant firing found during spontaneous or evoked activity, we performed whole cell patch-clamp recordings of PC from the Crus I/II area in sagittal cerebellar slices. No differences were found in passive membrane properties (resting potential, input resistance, membrane capacitance) between WT and *Cntnap2* KO mice (**Figure 3A**). Similarly, no differences in action potential (AP) threshold were found between genotypes (**Figure 3B**). The injected current needed to evoke the first action potential (rheobase) was smaller in *Cntnap2* KOs (**Figure 3C**), suggesting increased excitability. To specifically study PC intrinsic excitability, we measured the neuronal firing frequency elicited by depolarizing current steps of increasing amplitude. Both genotypes showed a linear current-frequency increase, as described for PC (F. R. Fernandez, Engbers, & Turner, 2007; Llinas & Sugimori, 1980) but, as expected, PC in *Cntnap2* KO mice fired at a higher frequency for the same injected current, indicating higher excitability (**Figure 3D**).

**Figure 3.**
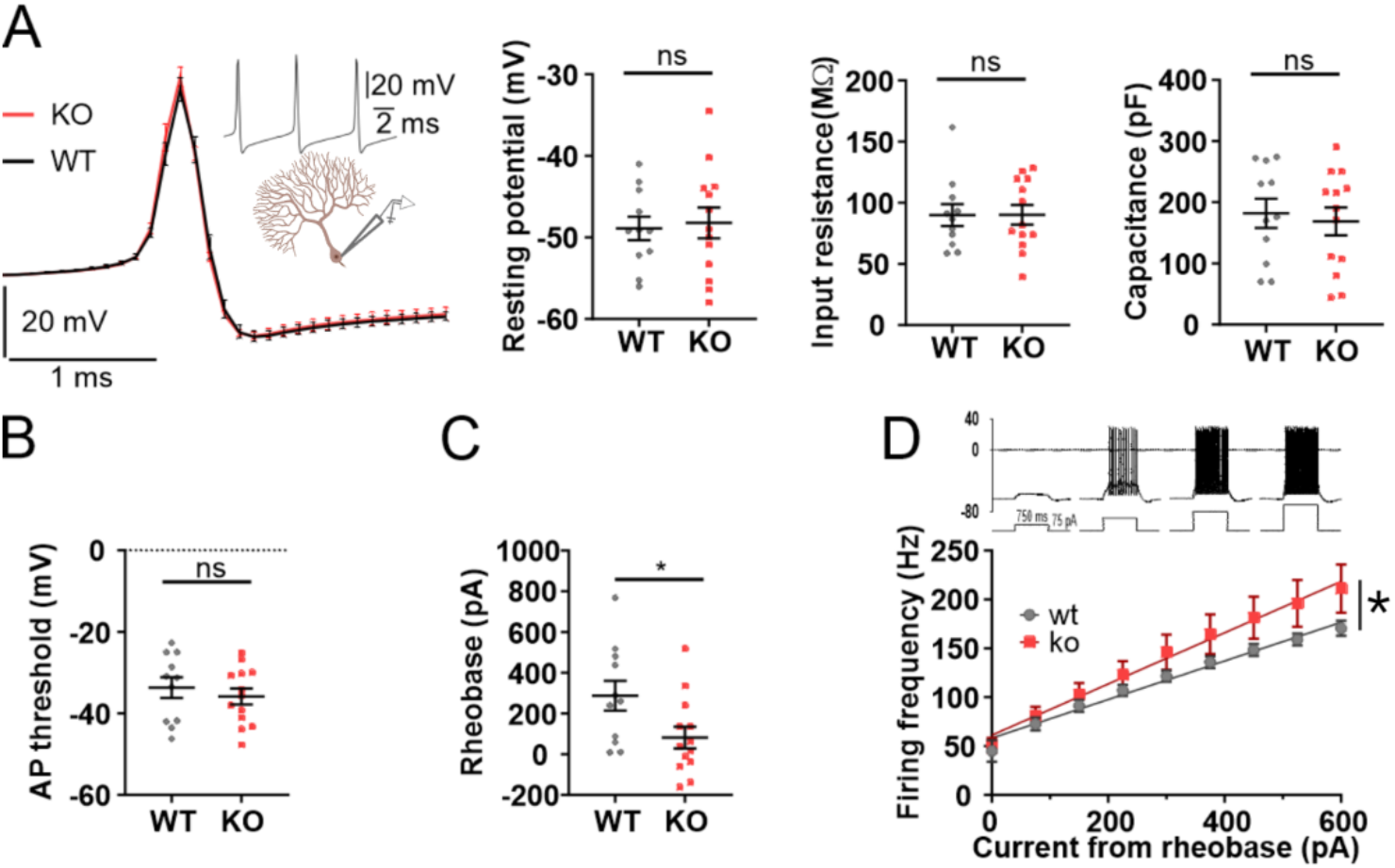
Increased intrinsic excitability of Purkinje cells in *Cntnap2* KOs. (A) Representative action potential from WT and KO mice. No statistically significant differences were found in passive membrane properties between WT and KO PC (resting potential, input resistance, capacitance). Student *t* test, p>0.05 (B) Action potential threshold. (C) Rheobase. Student *t* test, *p<0.05. (D) Intrinsic excitability as a measurement of firing frequency upon current step increases. Two Way ANOVA mixed effect (interaction genotype x current *p<0.05). All data are presented as mean ± S.E.M. n=11-13 neurons/genotype.

### Reduced dendritic complexity of Purkinje cells in *Cntnap2* mice

Recently, *Cntnap2* has been reported to modulate the development of PC (Argent et al., 2020). Further, we previously demonstrated that *Cntnap2* KO mice show defects in spine stabilization in the cerebral cortex (Gdalyahu et al., 2015). To determine whether the electrophysiological alterations observed in PC in *Cntnap2* KOs are associated with morphological changes, we characterized PC morphology in the patched neurons by biocytin staining (**Figure 4A**). We found that the overall length of the cells is significantly smaller in *Cntnap2* mice (**Figure 4B**). Sholl analysis of dendritic complexity demonstrated that PC of KO mice not only are smaller but also have a less complex dendritic arbor (**Figure 4C**). Last, no statistically significant differences in spine density were found between genotypes (**Figure 4D**).

**Figure 4.**
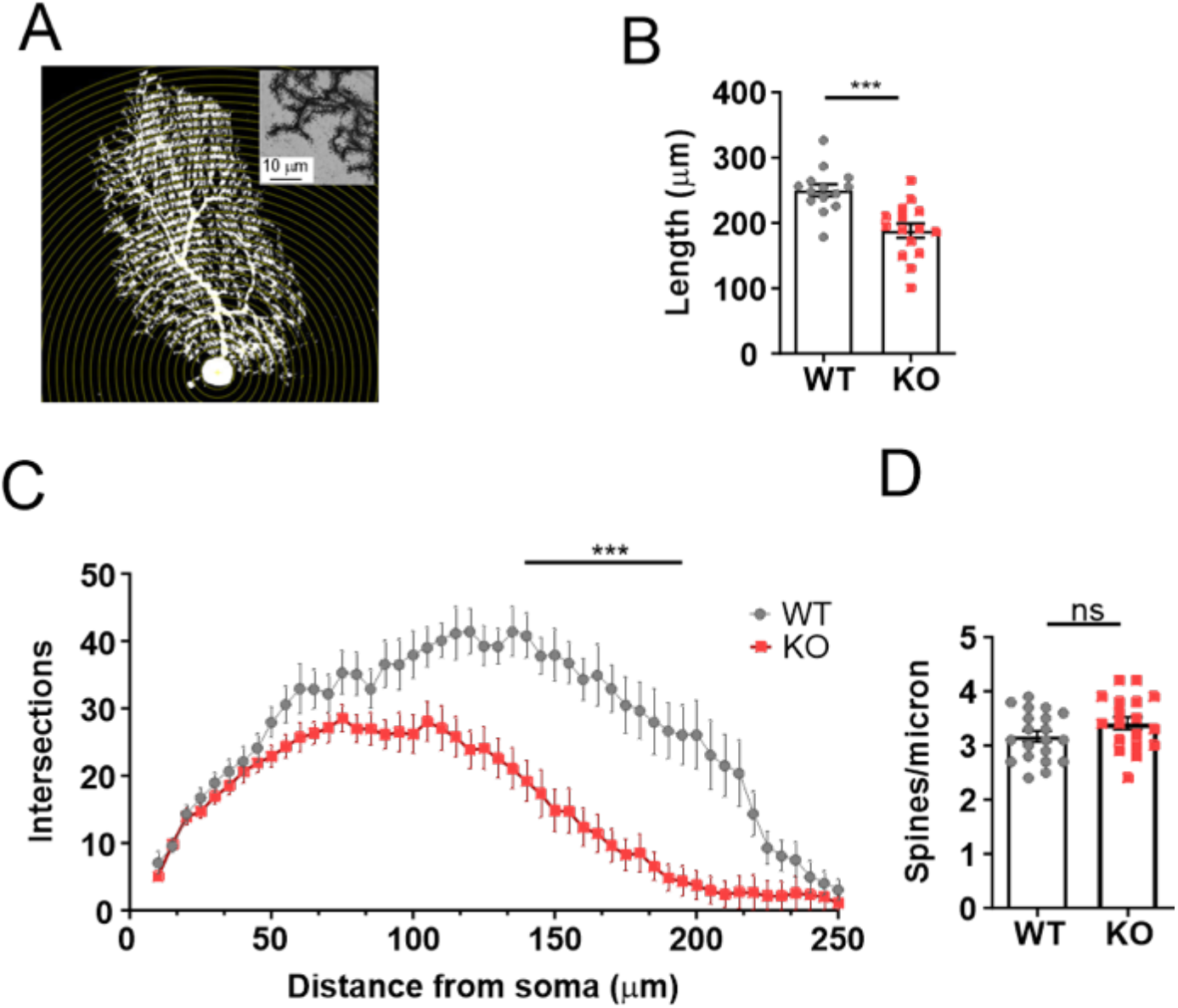
Neuroanatomical characterization of Purkinje cells. (A) Representation of Sholl analysis applied to a Purkinje cell. (B) Purkinje cell length from soma to the most apical point. Student’s t-test, p<0.001 (C) Number of intersections as a function of their distance from the soma, representing the complexity of arborization. Two-Way ANOVA with repeated measures. (D) Spine density. Data are presented as mean and ± S.E.M. *p<0.05, **p<0.01, ***p<0.001.

## Discussion

There is growing evidence for the involvement of the cerebellum in ASD through its role in integrating sensory neural signals. In this work, we investigated cerebellar integration of sensory information *in vivo* in the *Cntnap2* mouse model of autism, an autism-linked gene involved in cerebellar development and function. We used a well-characterized paradigm to study cerebellar integration of sensory function, where the evoked activity of PC, the sole output of the cerebellum, is analyzed upon whisker stimulation (Cheron, Dan, & Marquez-Ruiz, 2013). Whisker-related sensory information is conveyed to the cerebellar area Crus I/II, which has been widely associated with ASD (M. Fernandez, Sierra-Arregui, & Penagarikano, 2019), making it an ideal area of study. We found that the evoked response of PC to sensory stimuli (electrical stimulation of the whisker pad) was strikingly different between WT and *Cntnap2* KO mice, as denoted by the generated SEP. Whisker information reaches the cerebellum via two routes, the pontine nuclei-parallel fiber pathway and the inferior olive-climbing fiber pathway (Kleinfeld, Berg, & O’Connor, 1999), generating the two distinct types of firing that characterize PC: SS and CS, respectively. Individual analysis of each type of spike upon stimulation revealed that the timing of appearance of the stimulus-evoked CS, anticipated in *Cntnap2* KOs, was driving the observed differences in SEP, indicating an altered integration of climbing fiber inputs. In agreement with this, dysfunction of the olivocerebellar circuit in *Cntnap2* mice was proposed as responsible for the decreased response probability shown by this model in the eye-blink conditioning test, another paradigm widely used to study cerebellar function (Kloth et al., 2015). The appearance of the stimulus-evoked CS has been shown to be modulated by activity from the somatosensory cortex, since the suppression of S1 activity abolishes CS appearance upon tactile stimulation (Shimuta, Sugihara, & Ishikawa, 2020). Further, diminishing S1 activity lengthens and enhancing it shortens CS appearance latency (Brown & Bower, 2002). These data would suggest that increased activity of S1 could be responsible for the observed CS anticipation in the *Cntnap2* model. In fact, decreased inhibitory markers and increased excitation/inhibition ratio leading to higher cortical sensory gain have been described in *Cntnap2* mice (Antoine, Langberg, Schnepel, & Feldman, 2019; Penagarikano et al., 2011), which could account for the observed deficits.

CS have been recently shown to control the information encoded by SS activity (Streng, Popa, & Ebner, 2017) in a way that a dynamic relationship between CS and SS is necessary for the initiation of sensory-driven behavior (Tsutsumi et al., 2020). We observed reduced spontaneous CS firing frequency as well as increased irregularity in the SS timing, despite normal SS firing frequency. The rhythmicity of PC inter-spike intervals is thought to affect the transmission of information to downstream neurons. It is worth noting that increased coefficient of variation has been reported in other animal models of autism with cerebellar dysfunction including *CACNA1A* (Damaj et al., 2015; Jayabal, Chang, Cullen, & Watt, 2016), *CAMK2B* (Kury et al., 2017; van Woerden et al., 2009) and *KCNMA1* (Chen et al., 2010; Laumonnier et al., 2006), being the last two also associated with a reduced CS firing frequency. CS are originated in the dendritic tree of PC (Davie, Clark, & Hausser, 2008). In the adult stage, each PC is innervated by a single climbing fiber that presents hundreds of release sites depolarizing the bulk of the dendritic tree to induce CS activity. Although spine density is similar, the reduced PC arborization found in *Cntnap2* mice imply that each neuron’s total number of synapses is substantially decreased and could potentially be responsible for the diminished spontaneous CS frequency. Given this reduced CS firing, the observation of increased PC excitability in *Cntnap2* KOs was unexpected. One possible explanation is that dendritic input from CF is not represented by somatic current injection. PC somatic excitability has been shown to be regulated by small-conductance potassium (SK) channels, as SK blockade leads to increased neuronal firing (Grasselli et al., 2020). CNTNAP2 is known to cluster voltage-gated potassium channels, including KCNA1 (Kv1.1), KCNA2 (Kv1.2) (Poliak et al., 1999). Reduced expression of such channels was observed in hippocampal tissue resected from epileptic patients harboring *CNTNAP2* mutations (Strauss et al., 2006). Whether the increased PC excitability is due to reduced expression or mislocalization of potassium channels and its contribution to circuit dysfunction remains to be investigated.

In summary, we report an altered cerebellar response to evoked sensory stimuli in alert *Cntnap2* mice. This alteration is correlated with neuroanatomical and electrophysiological dysfunction of PC. This mouse model, therefore, provides a valuable tool to study the basis for the altered integration of sensory information observed in ASD.

## Materials and methods

### Animals

Adult (8-10 weeks) male mutant mice lacking the *Cntnap2* gene (Cntnap2-KO) and age-matched wild-type controls (C57BL/6 background) were purchased from The Jackson Laboratory (Bar Harbor, ME, USA). Animals were housed 4-5 per cage on a 12-12 h light/dark cycle, at 21-23°C and 65-70% humidity. Food and water were provided *ad libitum*. Animal maintenance and experimental procedures were executed following the guidelines of animal care established by the European Communities Council Directive 2010/63/EU, as well as in agreement with the Spanish Legislation (Royal Decree 53/2013). Procedures were also approved by the Ethics Committee for Animal Welfare (CEBA) of the University of the Basque Country (UPV/EHU) and the Pablo de Olavide University (UPO).

### *In vivo* electrophysiology

We followed the procedures described in Sánchez-León et al. (Sanchez-Leon et al., 2021). Briefly, stereotaxic surgery was performed to open a craniotomy (2mm ø) following the Allen Brain Atlas coordinates for the right Crus I/II area (AP: −6,6 mm; and L: −2,6 mm, relative to Bregma). During surgery, two small bolts were cemented in the skull to immobilize the head during the recording sessions, and a silver reference electrode was placed on the surface of the parietal cortex. The surface of the craniotomy was protected with bone wax (Ethicon, Johnson & Johnson) until recording sessions. After the surgery, mice were allowed to recover for at least two days. For *in vivo* recordings, the animal’s head was fixed to the recording setup, consisting of a treadmill with an infrared sensor for monitoring locomotor activity. All experiments were carried out with an amplifier (BVC-700A, Dagan corporation, MN, USA) connected to a dual extracellular-intracellular headstage (8024 Dual Intracellular & Extracellular Headstage, MN, USA). For SEP recordings we used a micropipette with a tip diameter between 8-10 μm, subsequently the pipette was filled with 3 M NaCl and placed on a micromanipulator (Narishige MO-10, Japan). Whisker stimulation was performed with a pair of flexible steel electrodes (Strand ø: 50.8 μm; Coated ø: 228.6 μm; Multi-Stranded PFA-Coated Stainless Steel Wire, A-M Systems, WA, USA) inserter under the skin of the right whisker pad. The electrical stimulus consisted of a single square pulse (0.2 ms; 0.5-1 mA) delivered by an isolation unit (Cibertec ISU 210 BIP) connected to a stimulator device (CS420, Cibertec), applied every 10±2 s. For single-Purkinje cell activity, a micropipette with a tip diameter around 1-2 μm was filled with 3 M NaCl and placed on a micromanipulator (Narishige MO-10, Japan). The pipette was inserted in the area of interest at ~2 μm/s and spikes were detected based on visual (2002C and 2004C, Tektronix, OR., USA) and auditory cues (Audio monitor 3300, A-M Systems, WA., USA).

Data were collected with a CED micro1401-3 data acquisition unit and sampled at 25 kHz. SEP analysis was performed with EEGLAB rev.14.1.2 toolbox using the Matlab 2015a software package. Recorded data were segmented into 70 ms windows using the electrical stimulation as trigger and baseline was corrected by subtracting the mean voltage level in the first 20 ms interval of the window (before whisker stimulus). Data were averaged for each genotype to obtain the average SEP and temporal periods were statistically compared. For single-cell recording analysis, only well isolated neurons recorded during at least 100s were considered. A DC remove process (time constant (s): 0,001-0,0004) was applied to reduce DC level drifts, and spikes were detected based on threshold-crossing algorithm of Spike2 software. All spikes were visually confirmed and Purkinje cells were identified by the presence of CS. Subsequently, SS and CS of each neuron were analyzed using a Matlab custom-made script. The Predominant Firing Rate represent the mode of the firing rate, the Frequency of the firing rate that appears most often. The Coefficient of variation CV and CV2, as a measure of firing regularity, were calculated following the formulas described in Holt et al. (Holt, Softky, Koch, & Douglas, 1996) 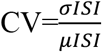, where ISI represents the inter-spike interval. 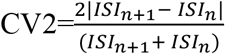. The Complex Spikes were sorted manually offline for the duration of the recording.

### *Ex vivo* electrophysiology

Mice were anesthetized with isoflurane and decapitated. The brain was rapidly submerged in icecold cutting solution containing (in mM): 20 NaCl, 2.5 KCl, 0.5 CaCl_2_, 7 MgCl_2_, 1.25 NaH_2_PO_4_, 85 sucrose, 25 D-glucose and 60 NaHCO_3_, saturated with 95% O_2_/5% CO_2_ (carbogen). The cerebellum was separated from the rest of the brain and the right part cut and glued to the cutting-dish. Parasagittal slices (250 μm thick) containing the Crus I area were prepared using a Campden Ci 7000smz-2 vibroslicer. Immediately upon cutting, slices were submerged in an artificial cerebrospinal fluid (aCSF) containing (in mM): 126 NaCl, 2.5 KCl, 1.2 MgCl_2_, 2.4 CaCl_2_, 1.2 NaH_2_PO_4_, 11.1 D-glucose, 21.4 NaHCO_3_, 0.1 ascorbic acid and 0.4 kynurenic acid, bubbled with carbogen at room temperature (22-24 °C), and left to recover for at least 1 h. Slices were constantly perfused with aCSF at 33°C. PCs were visualized using infrared differential interference contrast light (Olympus BX52WI). Patch pipettes pulled from thin borosilicate capillary glass (World Precision Instruments) with a Sutter P-97 horizontal puller had a resistance of 3–5 MΩ. Internal solution contained (in mM): 115 K-gluconate, 10 HEPES, 11 EGTA, 2 MgCl_2_, 10 NaCl, 2 MgATP, 0.25 Na_2_GTP and biocytin (5 mg/ml, B4261 Sigma-Aldrich), pH 7.3, adjusted with NaOH; osmolality ± 275 mOsm.

Intrinsic properties were studied in whole-cell voltage-clamp and in current-clamp mode by injecting a hyperpolarizing bias current (less than −500 pA) to hold membrane potential between −60 and −65 mV and keep the neuron silenced during the rheobase study. Membrane potentials were not corrected for the liquid junction potential between intra and external solution (−12.7 mV). Neurons in which holding current was greater than −500 pA and experiments in which access resistance was higher than 16 MΩ were discarded and not included for analysis. Signals from the patch pipette were recorded with a MultiClamp 700B amplifier, digitized at 10–20 kHz and low-pass filtered at 2–5 kHz with a Digidata 1440A analog-to-digital converter and analyzed off-line using Clampfit 10.7 software (Molecular Devices). Intrinsic excitability was determined in response to increasing depolarizing current pulses (+75 pA) of 750 ms duration injected from hyperpolarized holding currents. Rheobase was registered for each neuron as the net depolarizing current capable of inducing the first action potential (AP). PCs have been described to have different integrative properties making that neurons with similar resistances tend to respond with different firing frequencies to current pulses of similar amplitude and duration (Llinas & Sugimori, 1980). In order to facilitate the comparison between firing frequency curves, rheobase was normalized to zero and only the instantaneous frequencies between the first six APs were analyzed. AP threshold for each neuron was calculated for the first spike as the voltage where dV/dt reaches 5% of the AP maximal rise slope.

### Neuroanatomy

PCs were filled during patch-clamp recordings with biocytin (Sigma) via passive diffusion. Filled slices were fixed in 4% paraformaldehyde in PBS for 24h at 4°C and incubated with Alexa 555/488-Streptavidin (1/500) for 48h at 4°C. Subsequently the slices were washed with PBS and mounted using Prolong Antifade Gold mounting medium (Invitrogen).

Super-resolution images were acquired using a confocal LSM880 Fast Airyscan microscope using a 25x/0.8 water (for dendrite analysis) and a 63x/0.4 oil (voxel size 49 × 49 × 211 nm, for spine analysis) Plan-Apochromat objectives. Sholl analysis and spine quantification were performed using Fiji imageJ software. For Sholl, the number of intersections of the dendritic arbour with concentric circles drawn at 5 μm intervals from the soma was counted. Spines were manually counted along a 10 μm distal dendrite.

### Statistical analysis

Data are displayed through the graphs as mean ± S.E.M. Statistical analyses were performed using Matlab (Math Works Inc.) and Prism (GraphPad). For comparisons between groups a one-way ANOVA or two-way ANOVA with repeated measures, when appropriate, were used, as indicated in each figure legend. The significance threshold was set at p=0.05 (ns=not significant, *p<0.05,**p<0.01, ***p<0.001). Figures were created with Biorender.

## Acknowledgments

The authors thank the SGIker Microscopy Core (UPV/EHU, ERDF, ESF) for technical support and Drs. Jorge Valero and Jan Tønnesen, from Achucarro Basque Center for Neuroscience, for help with neuroanatomical analyses. This work was supported by MCIU/AEl/FEDER, UE grant RTI2018-101427-B-I00 to OP, ERANET-NEURON grant nEUrotalk to OP and UPV/EHU grant GIU18/094 to OP; Israel Science Foundation (536/19) to SK; Spanish Ministry of Science (Europa Excelencia 15/02, SAF2016-78071-R to SK; BFU2017-89615-P from the Spanish MINECO-FEDER to JMR. MF holds a MINECO predoctoral fellowship (BES-2016-078420) and TS-A is a Basque Government predoctoral fellow (PRE-2020-2-0109).

## Competing interests

The authors declare no competing interests.

## Notes

### Competing Interest Statement

The authors have declared no competing interest.

## References

Antoine, M. W., Langberg, T., Schnepel, P., & Feldman, D. E. (2019). Increased Excitation-Inhibition Ratio Stabilizes Synapse and Circuit Excitability in Four Autism Mouse Models. Neuron, 101(4), 648–661 e644. doi:10.1016/j.neuron.2018.12.026

APA. (2013). Diagnostic and Statistical Manual of Mental Disorders, 5th Edition: DSM-5. American Psychiatric Association.

Argent, L., Winter, F., Prickett, I., Carrasquero-Ordaz, M., Olsen, A. L., Kramer, H.,… Becker, E. B. E. (2020). Caspr2 interacts with type 1 inositol 1,4,5-trisphosphate receptor in the developing cerebellum and regulates Purkinje cell morphology. J Biol Chem, 295(36), 12716–12726. doi:10.1074/jbc.RA120.012655

Becker, E. B., Zuliani, L., Pettingill, R., Lang, B., Waters, P., Dulneva, A.,… Vincent, A. (2012). Contactin-associated protein-2 antibodies in non-paraneoplastic cerebellar ataxia. J Neurol Neurosurg Psychiatry, 83(4), 437–440. doi:10.1136/jnnp-2011-301506

Brown, I. E., & Bower, J. M. (2002). The influence of somatosensory cortex on climbing fiber responses in the lateral hemispheres of the rat cerebellum after peripheral tactile stimulation. J Neurosci, 22(15), 6819–6829. doi:20026696

Chen, X., Kovalchuk, Y., Adelsberger, H., Henning, H. A., Sausbier, M., Wietzorrek, G.,… Konnerth, A. (2010). Disruption of the olivo-cerebellar circuit by Purkinje neuron-specific ablation of BK channels. Proc Natl Acad Sci U S A, 107(27), 12323–12328. doi:10.1073/pnas.1001745107

Cheron, G., Dan, B., & Marquez-Ruiz, J. (2013). Translational approach to behavioral learning: lessons from cerebellar plasticity. Neural Plast, 2013, 853654. doi:10.1155/2013/853654

D’Mello, A. M., Moore, D. M., Crocetti, D., Mostofsky, S. H., & Stoodley, C. J. (2016). Cerebellar gray matter differentiates children with early language delay in autism. Autism Res, 9(11), 1191–1204. doi:10.1002/aur.1622

Damaj, L., Lupien-Meilleur, A., Lortie, A., Riou, E., Ospina, L. H., Gagnon, L.,… Rossignol, E. (2015). CACNA1A haploinsufficiency causes cognitive impairment, autism and epileptic encephalopathy with mild cerebellar symptoms. Eur J Hum Genet, 23(11), 1505–1512. doi:10.1038/ejhg.2015.21

Davie, J. T., Clark, B. A., & Hausser, M. (2008). The origin of the complex spike in cerebellar Purkinje cells. J Neurosci, 28(30), 7599–7609. doi:10.1523/JNEUROSCI.0559-08.2008

Dawes, J. M., Weir, G. A., Middleton, S. J., Patel, R., Chisholm, K. I., Pettingill, P.,… Bennett, D. L. (2018). Immune or Genetic-Mediated Disruption of CASPR2 Causes Pain Hypersensitivity Due to Enhanced Primary Afferent Excitability. Neuron, 97(4), 806–822 e810. doi:10.1016/j.neuron.2018.01.033

Ellegood, J., & Crawley, J. N. (2015). Behavioral and Neuroanatomical Phenotypes in Mouse Models of Autism. Neurotherapeutics, 12(3), 521–533. doi:10.1007/s13311-015-0360-z

Fernandez, F. R., Engbers, J. D., & Turner, R. W. (2007). Firing dynamics of cerebellar purkinje cells. J Neurophysiol, 98(1), 278–294. doi:10.1152/jn.00306.2007

Fernandez, M., Sierra-Arregui, T., & Penagarikano, O. (2019). Thew cerebellum and autism, more than motor control. In Behavioral Neuroscience. London, UK: IntechOpen.

Gao, J. H., Parsons, L. M., Bower, J. M., Xiong, J., Li, J., & Fox, P. T. (1996). Cerebellum implicated in sensory acquisition and discrimination rather than motor control. Science, 272(5261), 545–547. doi:10.1126/science.272.5261.545

Gdalyahu, A., Lazaro, M., Penagarikano, O., Golshani, P., Trachtenberg, J. T., & Geschwind, D. H. (2015). The Autism Related Protein Contactin-Associated Protein-Like 2 (CNTNAP2) Stabilizes New Spines: An In Vivo Mouse Study. PLoS One, 10(5), e0125633. doi:10.1371/journal.pone.0125633

Gordon, A., Salomon, D., Barak, N., Pen, Y., Tsoory, M., Kimchi, T., & Peles, E. (2016). Expression of Cntnap2 (Caspr2) in multiple levels of sensory systems. Mol Cell Neurosci, 70, 42–53. doi:10.1016/j.mcn.2015.11.012

Grasselli, G., Boele, H. J., Titley, H. K., Bradford, N., van Beers, L., Jay, L.,… Hansel, C. (2020). SK2 channels in cerebellar Purkinje cells contribute to excitability modulation in motor-learning-specific memory traces. PLoS Biol, 18(1), e3000596. doi:10.1371/journal.pbio.3000596

Holt, G. R., Softky, W. R., Koch, C., & Douglas, R. J. (1996). Comparison of discharge variability in vitro and in vivo in cat visual cortex neurons. J Neurophysiol, 75(5), 1806–1814. doi:10.1152/jn.1996.75.5.1806

Jayabal, S., Chang, H. H., Cullen, K. E., & Watt, A. J. (2016). 4-aminopyridine reverses ataxia and cerebellar firing deficiency in a mouse model of spinocerebellar ataxia type 6. Sci Rep, 6, 29489. doi:10.1038/srep29489

Kleinfeld, D., Berg, R. W., & O’Connor, S. M. (1999). Anatomical loops and their electrical dynamics in relation to whisking by rat. Somatosens Mot Res, 16(2), 69–88. doi:10.1080/08990229970528

Kloth, A. D., Badura, A., Li, A., Cherskov, A., Connolly, S. G., Giovannucci, A.,… Wang, S. S. (2015). Cerebellar associative sensory learning defects in five mouse autism models. Elife, 4, e06085. doi:10.7554/eLife.06085

Kury, S., van Woerden, G. M., Besnard, T., Proietti Onori, M., Latypova, X., Towne, M. C.,… Mercier, S. (2017). De Novo Mutations in Protein Kinase Genes CAMK2A and CAMK2B Cause Intellectual Disability. Am J Hum Genet, 101(5), 768–788. doi:10.1016/j.ajhg.2017.10.003

Laumonnier, F., Roger, S., Guerin, P., Molinari, F., M’Rad, R., Cahard, D.,… Briault, S. (2006). Association of a functional deficit of the BKCa channel, a synaptic regulator of neuronal excitability, with autism and mental retardation. Am J Psychiatry, 163(9), 1622–1629. doi:10.1176/ajp.2006.163.9.1622

Lidstone, D. E., Rochowiak, R., Mostofsky, S. H., & Nebel, M. B. (2021). A Data Driven Approach Reveals That Anomalous Motor System Connectivity is Associated With the Severity of Core Autism Symptoms. Autism Res. doi:10.1002/aur.2476

Llinas, R., & Sugimori, M. (1980). Electrophysiological properties of in vitro Purkinje cell somata in mammalian cerebellar slices. J Physiol, 305, 171–195. doi:10.1113/jphysiol.1980.sp013357

Marquez-Ruiz, J., & Cheron, G. (2012). Sensory stimulation-dependent plasticity in the cerebellar cortex of alert mice. PLoS One, 7(4), e36184. doi:10.1371/journal.pone.0036184

Melzer, N., Golombeck, K. S., Gross, C. C., Meuth, S. G., & Wiendl, H. (2012). Cytotoxic CD8+ T cells and CD138+ plasma cells prevail in cerebrospinal fluid in non-paraneoplastic cerebellar ataxia with contactin-associated protein-2 antibodies. J Neuroinflammation, 9, 160. doi:10.1186/1742-2094-9-160

Mostofi, A., Holtzman, T., Grout, A. S., Yeo, C. H., & Edgley, S. A. (2010). Electrophysiological localization of eyeblink-related microzones in rabbit cerebellar cortex. J Neurosci, 30(26), 8920–8934. doi:10.1523/JNEUROSCI.6117-09.2010

Penagarikano, O., Abrahams, B. S., Herman, E. I., Winden, K. D., Gdalyahu, A., Dong, H.,… Geschwind, D. H. (2011). Absence of CNTNAP2 leads to epilepsy, neuronal migration abnormalities, and core autism-related deficits. Cell, 147(1), 235–246. doi:10.1016/j.cell.2011.08.040

Poliak, S., Gollan, L., Martinez, R., Custer, A., Einheber, S., Salzer, J. L.,… Peles, E. (1999). Caspr2, a new member of the neurexin superfamily, is localized at the juxtaparanodes of myelinated axons and associates with K+ channels. Neuron, 24(4), 1037–1047. doi:10.1016/s0896-6273(00)81049-1

Proville, R. D., Spolidoro, M., Guyon, N., Dugue, G. P., Selimi, F., Isope, P.,… Lena, C. (2014). Cerebellum involvement in cortical sensorimotor circuits for the control of voluntary movements. Nat Neurosci, 17(9), 1233–1239. doi:10.1038/nn.3773

Rodenas-Cuadrado, P., Pietrafusa, N., Francavilla, T., La Neve, A., Striano, P., & Vernes, S. C. (2016). Characterisation of CASPR2 deficiency disorder--a syndrome involving autism, epilepsy and language impairment. BMC Med Genet, 17, 8. doi:10.1186/s12881-016-0272-8

Sanchez-Leon, C. A., Cordones, I., Ammann, C., Ausin, J. M., Gomez-Climent, M. A., Carretero-Guillen, A.,… Marquez-Ruiz, J. (2021). Immediate and after effects of transcranial direct-current stimulation in the mouse primary somatosensory cortex. Sci Rep, 11(1), 3123. doi:10.1038/s41598-021-82364-4

Scott, K. E., Schormans, A. L., Pacoli, K. Y., De Oliveira, C., Allman, B. L., & Schmid, S. (2018). Altered Auditory Processing, Filtering, and Reactivity in the Cntnap2 Knock-Out Rat Model for Neurodevelopmental Disorders. J Neurosci, 38(40), 8588–8604. doi:10.1523/JNEUROSCI.0759-18.2018

Shimuta, M., Sugihara, I., & Ishikawa, T. (2020). Multiple signals evoked by unisensory stimulation converge onto cerebellar granule and Purkinje cells in mice. Commun Biol, 3(1), 381. doi:10.1038/s42003-020-1110-2

Skefos, J., Cummings, C., Enzer, K., Holiday, J., Weed, K., Levy, E.,… Bauman, M. (2014). Regional alterations in purkinje cell density in patients with autism. PLoS One, 9(2), e81255. doi:10.1371/journal.pone.0081255

Strauss, K. A., Puffenberger, E. G., Huentelman, M. J., Gottlieb, S., Dobrin, S. E., Parod, J. M.,… Morton, D. H. (2006). Recessive symptomatic focal epilepsy and mutant contactin-associated protein-like 2. N Engl J Med, 354(13), 1370–1377. doi:10.1056/NEJMoa052773

Streng, M. L., Popa, L. S., & Ebner, T. J. (2017). Climbing Fibers Control Purkinje Cell Representations of Behavior. J Neurosci, 37(8), 1997–2009. doi:10.1523/JNEUROSCI.3163-16.2017

Tan, G. C., Doke, T. F., Ashburner, J., Wood, N. W., & Frackowiak, R. S. (2010). Normal variation in fronto-occipital circuitry and cerebellar structure with an autism-associated polymorphism of CNTNAP2. Neuroimage, 53(3), 1030–1042. doi:10.1016/j.neuroimage.2010.02.018

Tang, T., Xiao, J., Suh, C. Y., Burroughs, A., Cerminara, N. L., Jia, L.,… Lang, E. J. (2017). Heterogeneity of Purkinje cell simple spike-complex spike interactions: zebrin- and non-zebrin-related variations. J Physiol, 595(15), 5341–5357. doi:10.1113/JP274252

Tsutsumi, S., Chadney, O., Yiu, T. L., Baumler, E., Faraggiana, L., Beau, M., & Hausser, M. (2020). Purkinje Cell Activity Determines the Timing of Sensory-Evoked Motor Initiation. Cell Rep, 33(12), 108537. doi:10.1016/j.celrep.2020.108537

van Woerden, G. M., Hoebeek, F. E., Gao, Z., Nagaraja, R. Y., Hoogenraad, C. C., Kushner, S. A.,… Elgersma, Y. (2009). betaCaMKII controls the direction of plasticity at parallel fiber-Purkinje cell synapses. Nat Neurosci, 12(7’), 823–825. doi:10.1038/nn.2329

Wang, S. S., Kloth, A. D., & Badura, A. (2014). The cerebellum, sensitive periods, and autism. Neuron, 83(3), 518–532. doi:10.1016/j.neuron.2014.07.016

Wang, Y., Xu, Q., Zuo, C., Zhao, L., & Hao, L. (2020). Longitudinal Changes of Cerebellar Thickness in Autism Spectrum Disorder. Neurosci Lett, 728, 134949. doi:10.1016/j.neulet.2020.134949

